# Ultraviolet-A light increases mitochondrial anti-viral signaling protein in confluent human tracheal cells even at a distance from the light source

**DOI:** 10.1101/2021.05.11.443549

**Authors:** Gabriela Leite, Ali Rezaie, Ruchi Mathur, Gillian Barlow, Gil Y. Melmed, Mark Pimentel

## Abstract

Mitochondrial antiviral signaling (MAVS) protein mediates innate antiviral responses, including responses to certain coronaviruses such as severe acute respiratory syndrome coronavirus-2 (SARS-CoV-2). We have previously shown that ultraviolet-A (UVA) therapy can prevent virus-induced cell death in human ciliated tracheal epithelial cells (HTEpC) infected with coronavirus-229E, and that UVA treatment results in an increase in intracellular levels of MAVS. In this study, we set out to determine the mechanisms by which UVA light can activate MAVS, and whether local UVA light application can activate MAVS at locations distant from the light source (such as via cell-to-cell communication). MAVS levels were compared in HTEpC exposed to 2 mW/cm^2^ narrow band (NB)-UVA for 20 minutes and in unexposed controls, at 30-40% and at 100% confluency. MAVS levels were also compared in unexposed HTEpC treated with supernatants or lysates from UVA-exposed cells or from unexposed controls. Also, MAVS was assessed in different sections of confluent monolayer plates where only one section was exposed to NB-UVA. The results show that UVA increases the expression of MAVS protein. Cells in a confluent monolayer exposed to UVA were able to confer an elevation in MAVS in cells adjacent to the exposed section, and even cells in the most distant sections not exposed to UVA. In this study, human ciliated tracheal epithelial cells exposed to UVA demonstrate increased MAVS protein, and also appear to transmit this influence to distant confluent cells not exposed to light.

## Introduction

The human body has various defense mechanisms against infections, the most well-known of which involve innate immune responses where immune cells are recruited to sites of infection via cytokine signaling [1, 2]. Host intracellular responses to infection are also important, particularly in the defense against viruses. In the past decade, it has been discovered that mitochondria can mediate innate and adaptive immune responses via several mechanisms [3], including the production of mitochondrial anti-viral signaling (MAVS) protein [4].

The MAVS protein is primarily localized to the outer membrane of the mitochondria, and transduces signals from cytoplasmic retinoic acid-inducible gene I (RIG-I)-like receptors (RLRs) that recognize viral RNA [4]. Specifically, after recognition and binding of viral components, the RLRs RIG-I and melanoma differentiation-associated gene 5 (MDA5) interact with MAVS, activating transcription factors that induce expression of proinflammatory factors and antiviral genes [4]. However, some viruses have developed mechanisms to antagonize the activation of MAVS and evade this innate immune response. For example, the SARS-CoV-2 transmembrane glycoprotein M is thought to antagonize MAVS, thus impairing MAVS-mediated innate antiviral responses [5].

We recently showed that application of UVA light, under specific conditions, to human ciliated tracheal epithelial cells infected with CoV-229E, significantly improved cell viability and prevented virus-induced cell death, and that this was accompanied by decreases in the levels of CoV-229E spike (S) protein [6]. Moreover, cells treated with UVA light exhibited significantly increased levels of MAVS protein [6]. This suggested that UVA may activate MAVS. Further, in a first-in-human clinical trial in ventilated subjects with coronavirus disease 2019 (COVID-19), a 20-minute endotracheal UVA treatment daily for 5 days resulted in significantly decreased respiratory SARS-CoV-2 viral loads [7]. Interestingly, despite time-limited localized UVA therapy in this trial, average log_10_ changes in endotracheal viral load from baseline to day 6 was −3.2, suggesting a potential antiviral phenomenon beyond immediate localized effects.

In this study, we explore the effects of narrow band (NB)-UVA light on MAVS expression in uninfected human ciliated tracheal epithelial cells *in vitro*. We also explore whether the effects of UVA light were limited to cells directly exposed to UVA, or were also seen in cells not directly exposed to UVA.

## Materials and methods

### NB-UVA effects on MAVS

Primary human tracheal epithelial cells isolated from the surface epithelium of human trachea (HTEpC, lot n° 454Z019.11, PromoCell GmbH, Heidelberg, Germany) were cultured at 37°C (5% CO_2_) in 60×15mm standard tissue culture dishes (cat. 351007, Corning, NY, USA) with Airway Epithelial Cell Growth Medium (cat. C-21060, PromoCell) prepared with SupplementMix (cat. C-39165, PromoCell) and Gibco antibiotic-antimycotic solution (cat. 15240096, ThermoFisher Scientific, MA, USA).

Once the cells reached 10^5^ cells per plate (30-40% confluency), HTEpC were washed 3 times with sterile 1x PBS pH 7.4 (cat. 10010072, ThermoFisher), and fresh media was added to each plate. Cells were exposed to 2 mW/cm^2^ of NB-UVA for 20 minutes based on previously validated ideal UVA irradiation levels [6]. Unexposed cells were used as controls. After 24 hours the supernatants were collected, and cell were washed 3 times with sterile 1x PBS, pH 7.4. Following the removal of any remaining PBS, cells were lysed in the plate using 1 mL of RTL buffer from an AllPrep DNA/RNA/Protein isolation kit (Qiagen, Hilden, Germany). Experiments were performed in triplicate.

### NB-UVA effects on MAVS signal transmission to unexposed UVA- naïve cells

To determine whether the activation of MAVS caused by exposure to NB-UVA light could be transmitted to naïve, unexposed HTEpC, and to begin to elucidate the mechanisms involved, three experiments were performed:

- To determine if an extracellular mediator was involved, supernatants from 30-40% confluent HTEpC that were exposed to NB-UVA were transferred to 30-40% confluent naïve HTEpC.
- To determine if an intracellular mediator was involved, cell lysates from 30-40% confluent HTEpC that were exposed to NB-UVA (after supernatant removal) were transferred to 30-40% confluent naïve HTEpC.
- To determine if cell-to-cell signaling was involved, areas of 100% confluent HTEpC exposed or not exposed to NB-UVA were analyzed.

### NB-UVA effects on MAVS signal transmission via extracellular mediators

Supernatants collected from UVA-exposed and control HTEpC from the previous experiment were transferred to a new 60×15mm tissue culture dish containing 10^5^ naïve HTEpC (i.e. cells that were never exposed to UVA). Before receiving the supernatant from UVA-exposed or control cells, the naïve HTEpC were washed 3 times with sterile 1x PBS, pH 7.4. The PBS was completely removed, and 4 mL of the supernatant collected from UVA-exposed or control HTEpC were added to the naïve cells. After 24 hours of incubation, the cells were washed 3 times, and were then lysed in the plate using 1 mL of RTL buffer from an AllPrep DNA/RNA/Protein isolation kit (Qiagen). Experiments were performed in triplicate.

### NB-UVA effects on MAVS signal transmission via intracellular mediators

HTEpC were cultured at 37°C (5% CO_2_) in 60×15mm standard tissue culture dishes (cat. 351007, Corning, NY, USA) with Airway Epithelial Cell Growth Medium (cat. C-21060, PromoCell) that included SupplementMix (cat. C-39165, PromoCell) and Gibco antibiotic-antimycotic solution (cat. 15240096, ThermoFisher Scientific, MA, USA).

Once the cells reached 10^5^ cells per plate (30-40% confluency), HTEpC were washed 3 times with sterile 1x PBS pH 7.4 (cat. 10010072, ThermoFisher), and fresh media was added to each plate. Cells were exposed to 2 mW/cm^2^ of NB-UVA for 20 minutes. Unexposed cells were used as controls. After 24 hours, the cells were washed 3 times with sterile 1x PBS, pH 7.4, scraped from the culture dishes, and transferred to a 15mL sterile tube. Cells were pelleted, and new fresh Airway Epithelial Cell Growth Medium was added. A single sterile 5 mm stainless steel bead (Qiagen) was added to each tube, and cells were lysed by vortexing the tube for 5 minutes. Lysates from UVA-exposed and control HTEpC were transferred to a new 60×15mm tissue culture dish containing 10^5^ naïve HTEpC (i.e. HTEpC that had never been exposed to UVA). Before receiving the lysate from UVA-exposed or control cells, naïve HTEpC were washed 3 times with sterile 1x PBS, pH 7.4. The PBS was completely removed, and 4 mL of the lysate from either UVA-exposed or control HTEpC were added to the naïve cells. After 24 hours of incubation, the cells were washed 3 times with sterile 1x PBS and were then lysed in the plate using 1 mL of RTL buffer from an AllPrep DNA/RNA/Protein isolation kit (Qiagen). Experiments were performed four times.

### NB-UVA effects on MAVS signal transmission via cell-to-cell signaling

HTEpC were cultured at 37°C (5% CO_2_) in 150mm dishes (cat. 430599, Corning) with Airway Epithelial Cell Growth Medium (cat. C-21060, PromoCell) prepared with SupplementMix (cat. C-39165, PromoCell) and Gibco antibiotic-antimycotic solution (cat. 15240096, ThermoFisher) until they reached 100% confluence.

On the day of NB-UVA therapy, cells were washed twice with sterile 1x PBS, pH 7.4, and fresh media was added. Each 150mm dish containing a 100% confluent monolayer of HTEpC was divided longitudinally into four sections, designated as areas 1, 2, 3 and 4, respectively (Fig 1). The NB-UVA emitting device was placed 2.3 cm from the bottom of the dish and approximately 2 mW/cm^2^ of NB-UVA was applied to area 1 for 20 minutes (S1 Fig). Experiments were performed four times.

**Fig 1.**
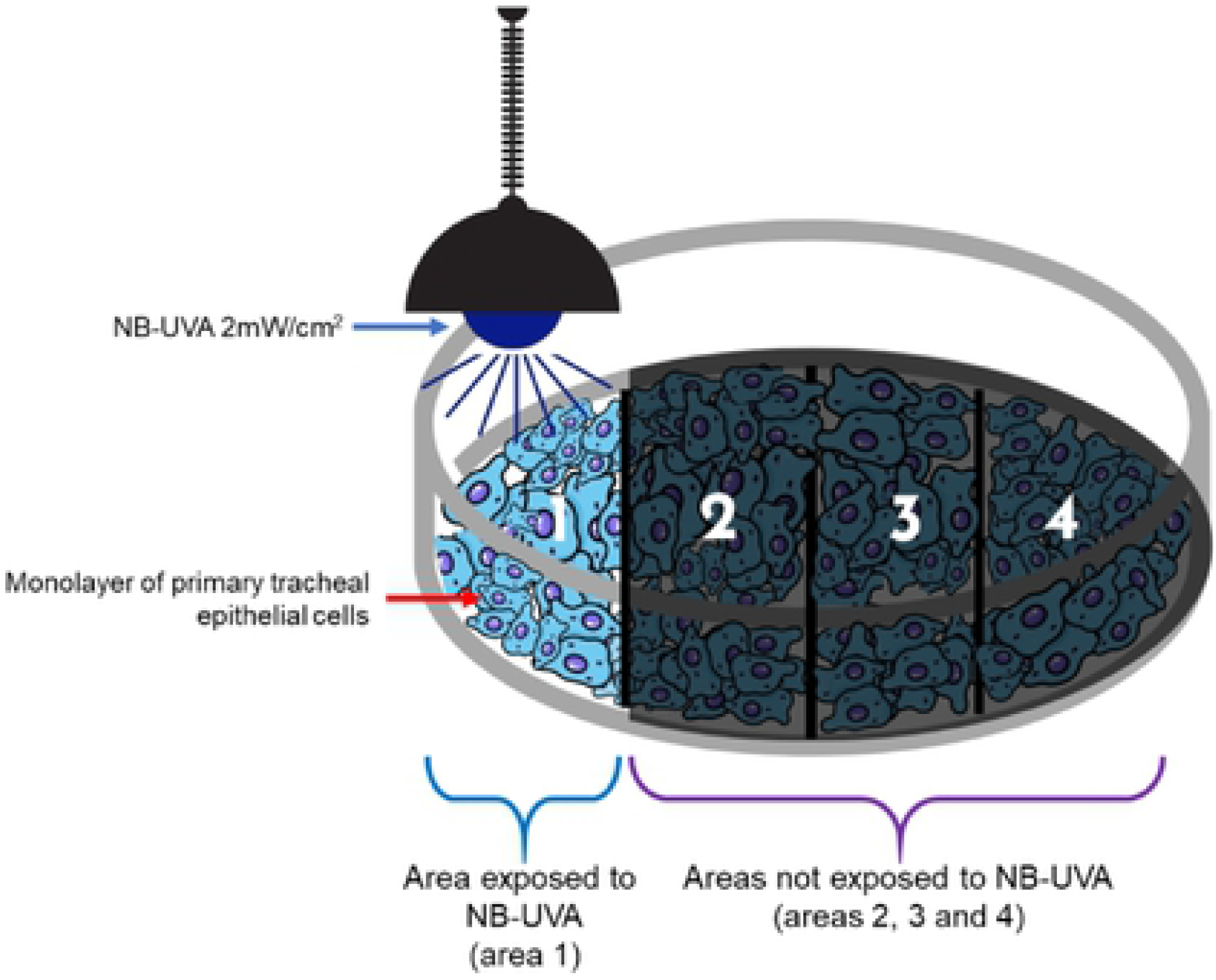
Schematic showing the design of experiments in which 100% confluent monolayer plates of primary tracheal epithelial cells (HTEpC) were partially exposed to 2 mW/cm^2^ NB-UVA for 20 minutes. NB-UVA was only applied to area 1. After UVA therapy, cells were collected from areas 4, 3, 2 and 1 in that order.

To prevent UVA leakage to other parts of the plate during therapy, areas 2, 3 and 4 were covered with a sterile barrier which blocked the passage of light through the top and sides of the plate (S1 Fig). During the course of the therapy NB-UVA intensity was constantly checked in unexposed areas (top, bottom, and sides) of the culture plates using a UV meter (SDL470, Extech, NH), to assure there was no UVA light in these areas (S1 Fig). UVA-treated plates were then re-incubated at 37°C (5% CO_2_) for 24h.

UVA-treated HTEpC plates were washed 3 times with sterile 1x PBS, pH 7.4, before harvesting the cells. 10 mL of sterile 1x PBS, pH 7.4, was added to the plate, and cells from area 4 were carefully scraped with a sterile Corning Cell Lifter (cat. 3008, Corning) and immediately transferred to a 15 mL sterile tube. Cells were pelleted at low speed (~1000 RPM) and lysed with one mL RTL buffer from an AllPrep DNA/RNA/Protein isolation kit (Qiagen).

The remaining UVA-exposed HTEpC from areas 1, 2 and 3 (still attached to the plate) were washed 3 times with sterile 1x PBS, pH 7.4. 10 mL of sterile 1x PBS, pH 7.4, was added to the plate and cells from area 3 were carefully scraped and lysed as described above. The same process was used to harvest the cells from areas 2 and 1 (in this order).

### Protein extraction and western blotting

AllPrep DNA/RNA/Protein Mini Kits (Qiagen) were used to extract total proteins from UVA-exposed and non-exposed HTEpC from all experiments, according to the manufacturer’s protocol. Total proteins were quantitated using Qubit Protein Assays (ThermoFisher) and equal loads of total protein were separated on a NuPAGE 4-12% Bis-Tris mini gel (NP0336BOX, ThermoFisher) and then transferred onto a Biotrace NT nitrocellulose membrane (27376-991, VWR). Total proteins were stained with Ponceau S solution (P7170, Sigma-Aldrich). The membrane was blocked with tris-buffered saline containing 3% bovine serum albumin (cat. A7030, Sigma-Aldrich) and 0.1% Tween 20 (P1379, Sigma-Aldrich) (TBS-T), and incubated overnight at 4°C with mouse anti-MAVS antibody (1:200; SC-166583, Santa Cruz Biotechnology) diluted in blocking solution. After washing in TBS-T, the membrane was then overlain with horseradish peroxidase (HRP)-conjugated goat anti-mouse IgG antibody (1:300; 5220-0286, SeraCare), washed in TBS-T, and exposed to enhanced chemiluminescence solution (RPN2235, GE Healthcare). Immunoreactive protein bands were imaged using an iBright FL1500 instrument (ThermoFisher) and analyzed using iBright Analysis software (ThermoFisher). Samples were normalized against total protein as determined from Ponceau S staining (MilliporeSigma, St. Louis, MO, US).

### Statistical Analysis

Graph construction and statistical analysis were performed with GraphPad Prism V. 9 (GraphPad Software, CA, USA). For all experiments, immunoreactive MAVS bands from nitrocellulose membranes were normalized against total protein (Ponceau S) before statistical analysis, using iBright Analysis software (ThermoFisher). MAVS relative densities (obtained after normalization) were compared between groups applying a non-paired t-test. Comparisons between each area from experiments with 100% confluent cell cultures were performed using paired t-test and ANOVA test. Significance level was set at p< 0.05.

## Results

### Narrow band-UVA (NB-UVA) increases MAVS protein levels in human non-confluent and confluent ciliated tracheal epithelial cells

Levels of MAVS were analyzed in primary tracheal epithelial cells (HTEpC) at 30-40% confluency which were exposed to 2 mW/cm^2^ NB-UVA for 20 minutes and in unexposed controls. Normalized MAVS levels, as detected by western blot, were increased in NB-UVA exposed cells when compared to unexposed controls (P=0.0193, Fig 2).

**Fig 2.**
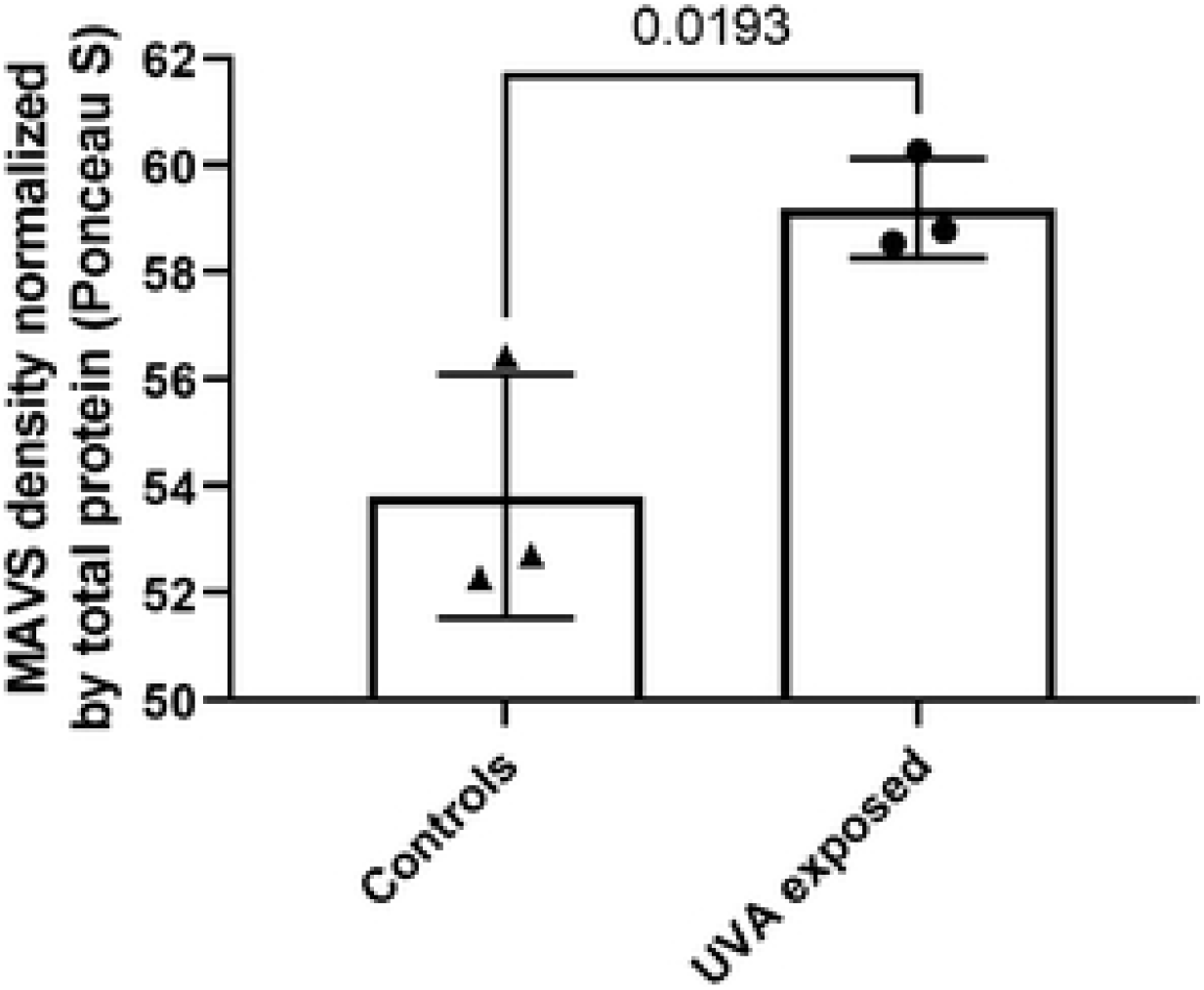
Normalized MAVS levels in 30-40% confluent HTEpC exposed to 2mW/cm^2^ NB-UVA for 20 minutes, and in unexposed controls.

In addition, when primary tracheal epithelial cells were grown in 100% confluent monolayers (as opposed to 30-40% confluency), normalized MAVS levels in area 1 were also significantly increased following exposure to 2 mW/cm^2^ NB-UVA for 20 minutes, when compared to levels in unexposed monolayers (P=0.0006, Fig 3, S2 Fig).

**Fig 3.**
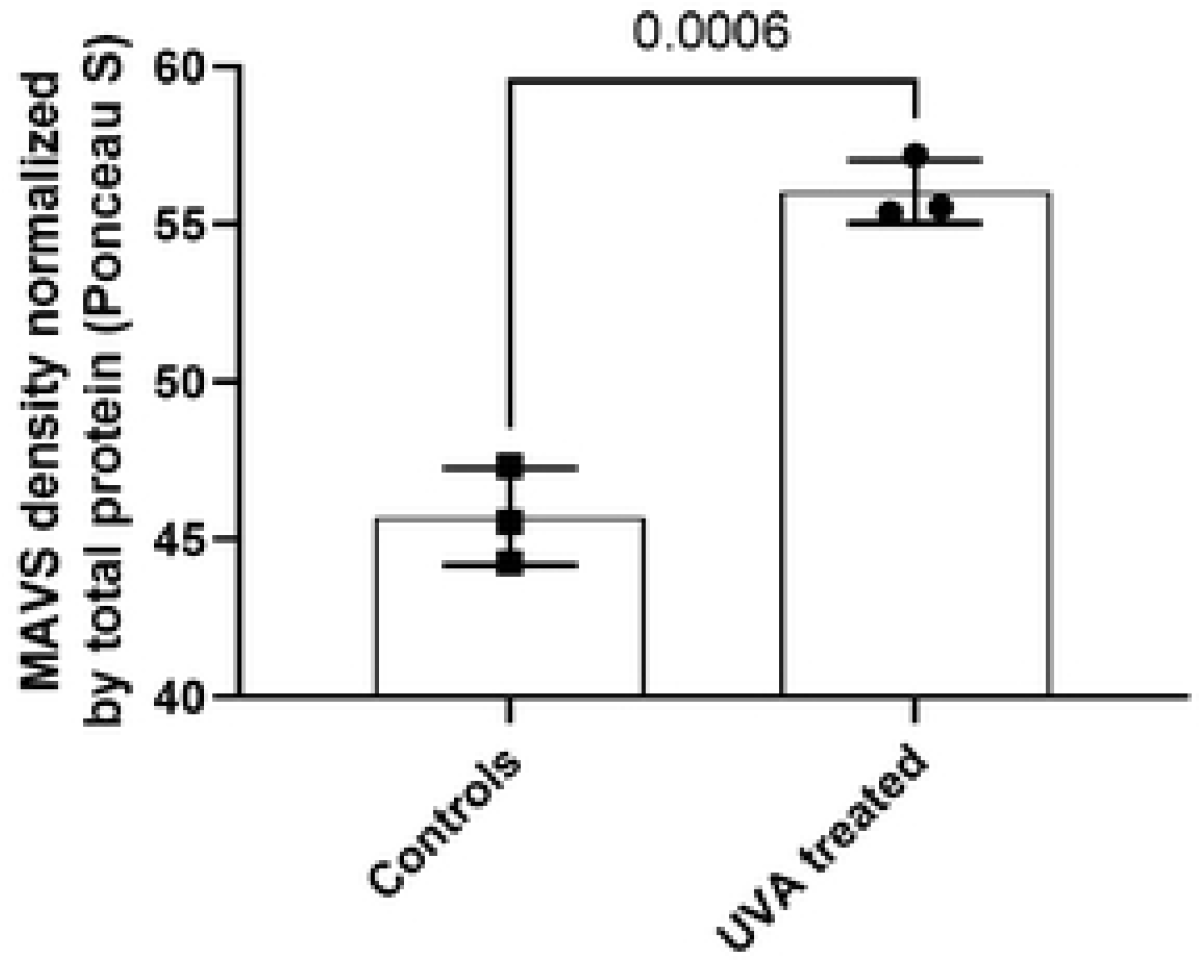
Normalized MAVS levels in 100% confluent HTEpC area 1 exposed to 2 mW/cm^2^ NB-UVA for 20 minutes, and in unexposed monolayer controls.

### MAVS is activated by cell-to-cell signaling after NB-UVA exposure

When naïve 30-40% confluent HTEpC were treated with supernatants from NB-UVA exposed 30-40% confluent HTEpC, no changes in MAVS levels were observed (P=0.4022, Fig 4). However, when naïve 30-40% confluent HTEpC were incubated with cell lysates from NB-UVA exposed 30-40% confluent HTEpC, normalized levels of MAVS tended to increase (Fig 5, P=0.1256).

**Fig 4.**
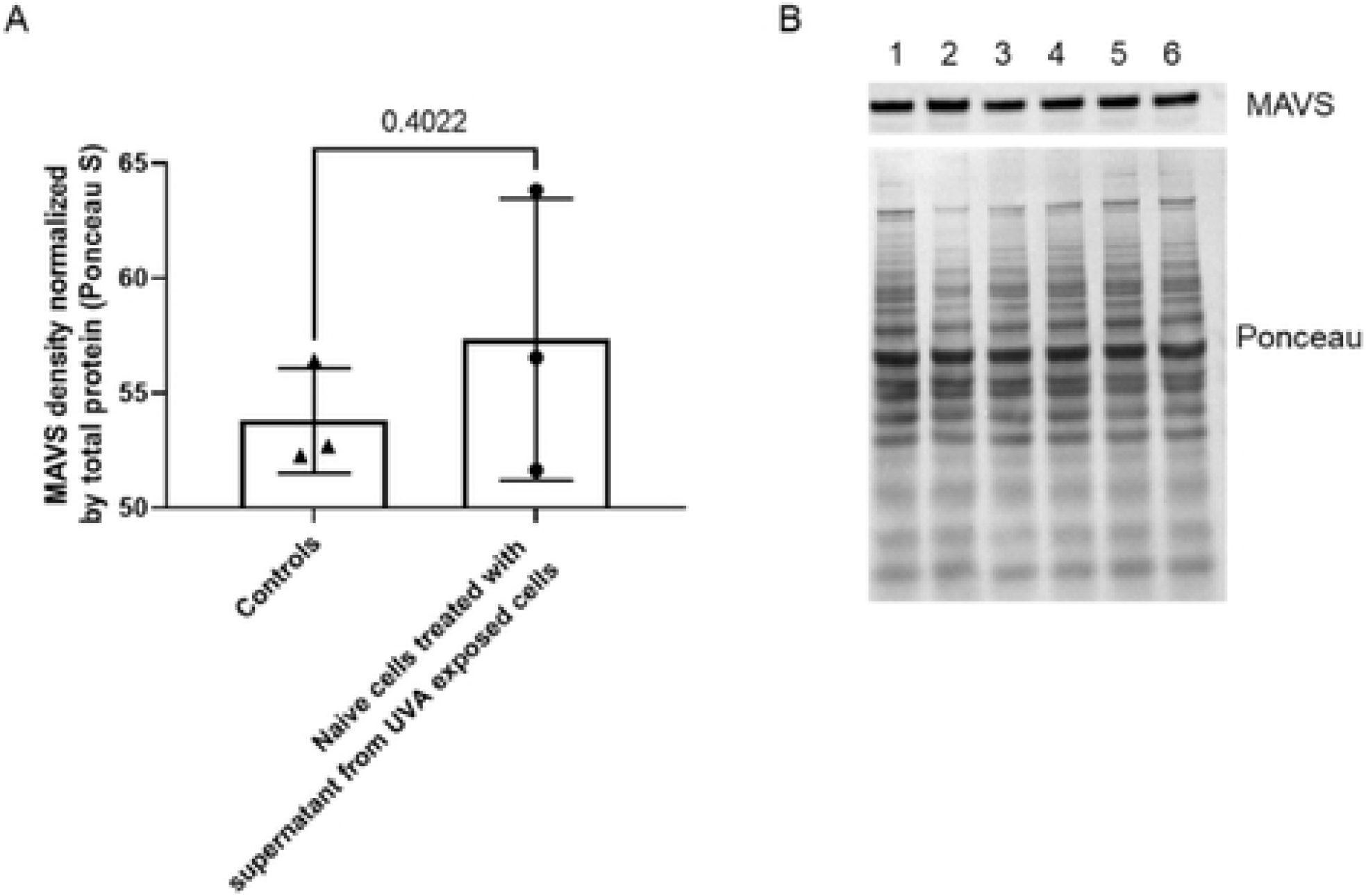
**A** – Normalized MAVS levels in 30-40% confluent naïve HTEpC treated with supernatants from 30-40% confluent NB-UVA exposed HTEpC, and in controls incubated with supernatants from unexposed 30-40% confluent HTEpC. **B** - Western blot of proteins extracted from 30-40% confluent naïve HTEpC treated with supernatant from 30-40% confluent NB-UVA exposed HTEpC (lanes 1, 2 and 3), and from controls treated with supernatant from 30-40% confluent unexposed HTEpC (lanes 4, 5 and 6).

**Fig 5.**
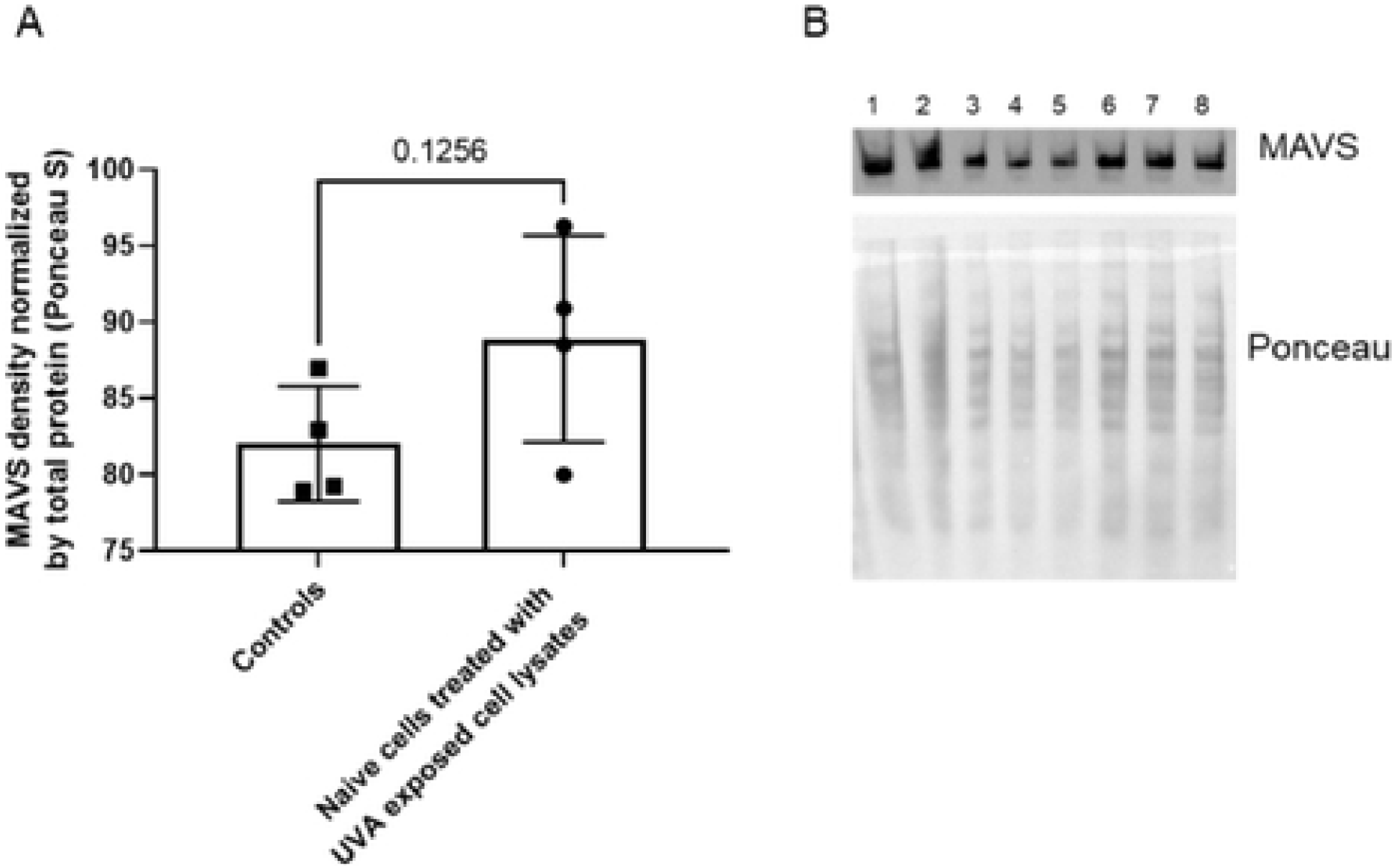
**A** –Normalized MAVS levels in 30-40% confluent naïve HTEpC treated with lysates from 30-40% confluent NB-UVA exposed HTEpC, and in controls incubated with lysates from 30-40% confluent unexposed HTEpC. **B** – Western blot prepared directly from lysates of 30-40% confluent naïve HTEpC incubated with lysates from 30-40% confluent NB-UVA exposed cells (lanes 1 to 4) and from lysates of controls incubated with lysates from 30-40% confluent unexposed HTEpC (lanes 5 to 8).

Next, levels of MAVS were analyzed in different areas of culture plates containing 100% confluent monolayers of HTEpC, after only one part of the plate (area 1) was exposed to 2 mW/cm^2^ NB-UVA for 20 min (Fig 1). Normalized MAVS levels gradually increased from area 4 (farthest unexposed area) through area 1 (exposed to NB-UVA) (ANOVA P=0.08, Fig 6A,B), and there was a statistically significant increase in MAVS levels in area 1 (exposed to NB-UVA) when compared to unexposed area 4 (P=0.0382, Fig 6A,B). Importantly, levels of MAVS were also significantly increased in unexposed areas 2 and 3 when compared to controls from unexposed plates (P=0.0289 and P=0.0402 respectively, Fig 6A). Normalized MAVS levels in area 4 (farthest unexposed area) also appeared to be higher than in controls, but did not reach statistical significance (P=0.1262, Fig 6A).

**Fig 6.**
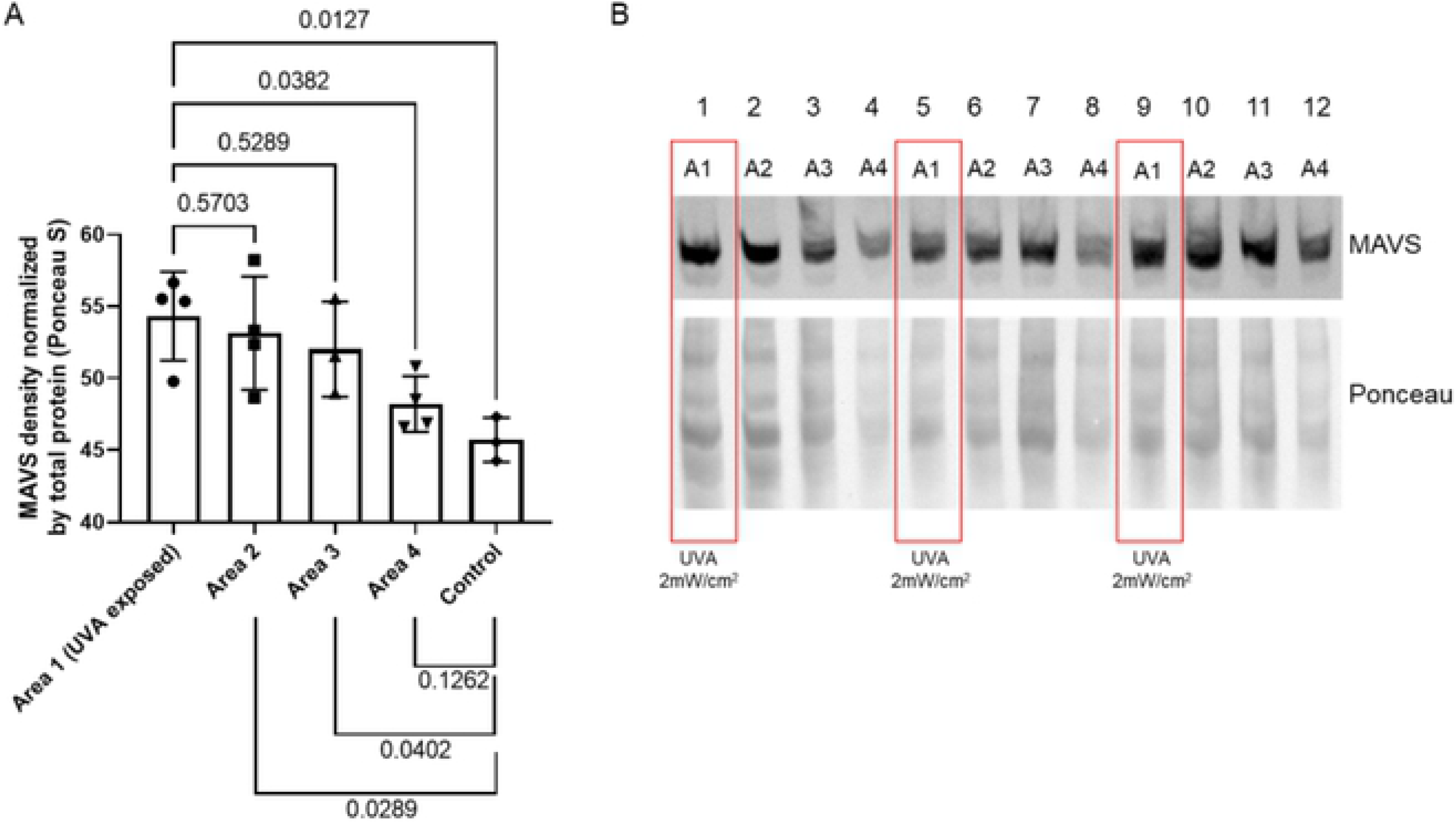
**A** – Normalized MAVS levels in 100% confluent HTEpC partially exposed to 2mW/cm^2^ NB-UVA for 20 minutes. Area 1 was directly exposed to NB-UVA, but areas 2, 3 and 4 were not exposed to NB-UVA. **B** –Western blot prepared from cell lysates of 100% confluent HTEpC from three experiments, exposed to NB-UVA (area 1 - lanes 1, 5 and 9) and from lysates of confluent HTEpC not exposed to NB-UVA from the same culture plate (area 2 - lanes 2, 6 and 10; area 3 - lanes 3, 7 and 11; area 4 - lanes 4, 8, and 12).

## Discussion

In this study, we show that narrow band UVA light increases the expression of the MAVS protein in uninfected human ciliated tracheal epithelial cells *in vitro*. In addition, in a confluent monolayer culture of these cells, the induction of MAVS protein is transmitted to cells not directly exposed to NB-UVA light. This transmission does not appear to be due to a secreted extracellular mediator, but more likely results from direct cell-to-cell signaling, and possibly a cytosolic mediator.

External UVA therapy has long been used in the treatment of skin conditions such as psoriasis, eczema and skin lymphoma, for which it is FDA-approved [8–11]. To explore the potential of internal UVA light therapy to treat microbial infections, we recently tested UVA efficacy against a variety of pathogens *in vitro*, and found that under controlled and monitored conditions, UVA light effectively reduced a variety of bacterial species (including *Klebsiella pneumoniae, Escherichia coli, Clostridioides difficile,* and others), the yeast *Candida albicans*, coxsackievirus group B, and coronavirus-229E [6]. Importantly, we found that human ciliated tracheal epithelial cells that were infected with coronavirus-229E and then treated with NB-UVA light *in vitro* exhibited increases in MAVS protein and survived infection [6]. These results suggested that the increased cell viability of coronavirus-229E-infected and UVA-treated cells, as compared to infected but untreated controls, might be due to activation of MAVS-mediated antiviral signaling pathways. In the present study, human ciliated tracheal epithelial cells were exposed to UVA light, without viral infection. The results confirmed that exposure to UVA light alone results in increased levels of the MAVS protein in these cells, demonstrating that this is a response to UVA light.

It is well recognized that the common cold, influenza and other viruses are seasonal and occur more often in winter and less in summer months. The mechanism for this is unclear, although data suggest that sunlight, and the production of vitamin D, may be important. Sunlight has historical importance in medicine – for example, during the H1N1 influenza pandemic of 1918–1919, it was suggested that the combination of access to sunlight and fresh air, together with strict hygiene and the use of face masks, may have lessened mortality among patients and staff at an ‘open-air’ hospital in Boston [12]. A systematic review of data regarding vitamin D levels and the current COVID-19 pandemic suggests that sunlight and elevated vitamin D levels may improve outcomes [13]. Although the trials selected for inclusion in the latter study had heterogenous results, these and other historical data [12] suggest that exposure to sunlight, and thus to UVA, may be beneficial in combating viral infections.

Under normal physiologic conditions, MAVS protein levels are low, due in part to binding of human antigen R as well as microRNAs to elements in the 3’UTR of the MAVS mRNA [4]. Following recognition and binding of viral components, the N-terminal caspase recruitment domains (CARDs) of RLRs are ubiquitinated and bind to the CARD of MAVS, leading to aggregation of MAVS and activation of proinflammatory cytokines and antiviral interferon genes [4]. However, viruses can also evade these pathways – for example, the membrane glycoprotein M of SARS-CoV-2, the virus which causes COVID-19 [14], can interact with MAVS and impair MAVS aggregation and activation of antiviral responses [5]. In our preclinical studies, tracheal cells that were infected with CoV-229E and treated with UVA light also exhibited decreases in CoV-229E spike protein [6], which suggested to us that UVA light might also be an effective treatment for SARS-CoV-2.

The primary site of SARS-CoV-2 infection is the ciliated epithelial cells, associated with downstream characteristic bilateral ground-glass opacities [15]. The acute respiratory viral infection and subsequent inflammatory responses can result in compromised pulmonary function [14, 16–18] and death [19]. Secondary bacterial and fungal infections are also common, with ventilator-associated pneumonia (VAP) occurring in 31% of mechanically ventilated patients [20]. To test the safety and efficacy of UVA light as a potential treatment for SARS-CoV-2, we developed a novel UVA light emitting diode (LED)-based catheter device which can be inserted into an endotracheal tube to deliver UVA light in critically ill COVID-19 subjects [7]. In the first human study of mechanically ventilated COVID-19 subjects, all of whom had World Health Organization (WHO) symptom severity scores of 9 at baseline (10 is death) [21], subjects who were treated with endotracheally-delivered UVA light (treated for 20 minutes daily for 5 days) exhibited an average log_10_ decrease in SARS-CoV-2 viral load of 3.2 (p<0.001) by day 6 of therapy in endotracheal aspirates, and these accelerated reductions in viral loads correlated with 30-day improvements in the WHO symptom severity scores [7]. Moreover, the scale of the improvements, despite the fact that only small portions of the trachea were exposed to UVA light, suggested the possibility that the antiviral effects of UVA light might not be confined to cells directly exposed to UVA, but might also be transmitted to neighboring cells.

To explore the potential mechanisms underlying this transmission, we first took supernatants from UVA-exposed cells and added them to fresh plates of cells that were never exposed to UVA light. No increase in MAVS protein levels were seen in these cells, indicating that a secreted extracellular mediator was not involved. Next, to explore whether a cytosolic mediator was involved, we lysed UVA-exposed cells and non-exposed controls and added the lysates to fresh plates of cells that were never exposed to UVA light. There was a trend towards an increase in MAVS protein levels in naïve HTEpC incubated with lysates from UVA-exposed cells, but this did not reach significance. In contrast, when we compared MAVS levels in confluent monolayers of HTEpC directly exposed to UVA light and in adjacent areas from the same plate that were blocked from UVA light, we found that MAVS was not only increased in cells in area 1 (directly exposed to UVA light), but was also increased in cells in the adjacent areas 2, 3, and 4 which were blocked from direct UVA light, in a gradient that decreased with increasing distance from UVA-exposed cells. These findings confirm that an increase in MAVS in response to UVA light can be transmitted from directly exposed cells to neighboring unexposed cells, and suggest that cell-to-cell signaling is involved, although further work is required to determine the mechanisms involved.

While this study is provocative, future studies are needed to explore the potential of UVA to enhance innate intracellular immunity to viruses. For example, as already stated, SARS-CoV-2 suppresses MAVS. Understanding the mechanisms by which UVA light overrides this suppression would be important. This could include damage to single-stranded viral RNA. In addition, the effects of this MAVS activation might be important to study with *in vivo* models. Limited data suggest that MAVS and resultant intracellular production of interferon α might attract circulating immune cellular response to attack infected cells [4]. Interestingly, in our previous *in vitro* study, CoV-229E caused precipitous cell death which was mitigated by UVA [6]. This increased cell survival suggests that perhaps MAVS is a cell salvage pathway. This is also supported by the first in human study of UVA in intubated critically ill subjects with COVID-19 [7]. Two patients underwent bronchoscopy after 5 days of UVA application. There was no macroscopic evidence of inflammation. Further studies are needed to explore these concepts not addressed in this current study.

In conclusion, this study begins to unravel the possible mechanisms by which UVA light could influence innate intracellular immunity. It appears that NB-UVA increases MAVS protein levels in human ciliated tracheal epithelial cells. This increase in MAVS protein appears to be transmissible to adjacent cells not directly exposed to UVA light. Further, our results suggest that MAVS signal transmission involves cell-to-cell communication, and possibly a cytosolic (but not a secreted extracellular) mediator. This finding may underlie the benefits of UVA seen *in vitro* and in human studies of critically ill patients with COVID-19. The findings could have wide-ranging implications for the treatment of SARS-CoV-2, other coronaviruses and other RNA respiratory viruses such as influenza. Further work is needed to determine if this mechanism is an important factor in the seasonality of specific respiratory viral illnesses.

## Acknowledgments

The authors thank Chandrima Chatterjee for her assistance in preparing the figures. We would also like to thank Ewan Seo for applying our ideas to paper in the early CAD drawings for the human UVA intra-tracheal device. In addition, the authors thank Frank Lee for his support of the MAST Program’s COVID-19 research. Finally, we would like to thank Aytu Biosciences for their commitment to examining this technology in critically ill human subjects.

## Supporting information

**S1 Fig.** Monolayer plates of primary tracheal epithelial cells (HTEpC) partially exposed to NB-UVA. No UVA light was detected in unexposed areas of the plate (i.e. areas 2, 3, and 4).

**S1 Fig.** Western blot of proteins extracted from 100% confluent HTEpC exposed to NB-UVA (lanes 1, 2 and 4), and 100% confluent HTEpC that were not exposed to NB-UVA (Lanes 5, 6 and 7). Lane 3 (exposed to NB-UVA) was discarded due to poor total protein magnification.

## Notes

### Competing Interest Statement

The authors have declared no competing interest.

